# Evaluation of In2Care® mosquito stations for suppression of the Australian backyard mosquito, *Aedes notoscriptus*

**DOI:** 10.1101/2023.03.20.533559

**Authors:** Véronique Paris, Nick Bell, Thomas L. Schmidt, Nancy M. Endersby-Harshman, Ary A. Hoffmann

## Abstract

*Aedes notoscriptus* (Skuse) is a container-inhabiting mosquito endemic to Australia that vectors arboviruses and is suspected to transmit *Mycobacterium ulcerans*, the cause of Buruli ulcer. We evaluated the effectiveness of the In2Care® station, which suppresses mosquito populations via the entomopathogenic fungus, *Beauveria bassiana*, and the insect growth regulator pyriproxyfen, the latter of which is autodisseminated among larval habitats by contaminated mosquitoes. A field trial was conducted using 110 In2Care® stations in a 50,000 m^2^ area and results were compared to four control areas that did not receive the treatment. Efficacy was evaluated by comparing egg counts and measuring larvicidal impact in surrounding breeding sites. Laboratory experiments validated the effect of *B. bassiana* on adult survival. Results of this field trial indicate that, six weeks after the In2Care® stations were deployed, treatment site ovitraps contained 43% fewer eggs than control site ovitraps, and 33% fewer eggs after ten weeks, suggesting that the In2Care® station was able to reduce the egg density of *Ae. notoscriptus*. Population reduction remained evident for up to three weeks after In2Care® stations were removed. Treatment site ovitraps had significantly fewer *Ae. notoscriptus* eclosing than control site ovitraps, confirming the pyriproxyfen autodissemination feature of the stations. An average reduction of 50% in adult eclosion was achieved. Exposure to *B. bassiana* resulted in four-times higher mortality among adult mosquitoes. Additionally, using fresh In2Care® nettings led to an 88% decrease in average survival compared to four-week-old nettings. The use of In2Care® stations has potential for suppressing *Ae. notoscriptus* egg density.

## 1 INTRODUCTION

Mosquitoes of the genus *Aedes* continue to pose a significant threat to human and animal health. While much research has been focused on the yellow fever mosquito (*Aedes aegypti* (L.)) and the tiger mosquito (*Aedes albopictus* (Skuse)), other container-inhabiting and human-biting *Aedes* species have received less attention, limiting information about effective control measures. One such species is *Ae. notoscriptus*, a container-inhabiting mosquito that is broadly distributed in its native range in mainland Australia and has become invasive in other regions, including the Torres Strait Islands, New Zealand, Papua New Guinea, New Caledonia, Indonesia (Dobrotworsky 1965, Lee et al. 1987, Sunahara and Mogi 2004), and the United States (specifically California) (Metzger et al. 2021). *Aedes notoscriptus* is a vector of human arboviruses, such as Ross River virus and Barmah Forest virus, (Doggett and Russell 1997), is suspected to play a significant role in the transmission of *Mycobacterium ulcerans* (the bacterium responsible for Buruli ulcer (BU)) (Wallace et al. 2017), and is also a primary vector of dog heartworm (Russell 1985).

*Aedes notoscriptus* has greater dispersal capacity than other container-inhabiting *Aedes* (Watson et al. 2000, Trewin et al. 2019, Paris et al. 2023), potentially making localised control efforts more challenging. A recent pilot study used gravid traps to target *Ae. notoscriptus* in the Mornington Peninsula, Australia, where it has a potential role in the transmission of Buruli ulcer (source: https://www.health.vic.gov.au/infectious-diseases/beating-buruli-in-victoria; Mee et al., unpublished). This study aimed to reduce local *Ae. notoscriptus* numbers, but no reduction in the number of *Ae. notoscriptus* eggs was observed. This outcome is likely in part due to the species’ ability for long-distance dispersal, as indicated by genetic data (Paris et al., 2023). In addition, the species has been shown to utilize cryptic water-filled containers for immature mosquito development (Montgomery and Ritchie 2002, Trewin et al. 2019) in properties that cannot be easily accessed to apply larvicides.

One potential solution for controlling pests with high dispersal rates in an environment with an abundance of cryptic breeding sites is through autodissemination of insecticides by the target pest. This method has been implemented in both agriculture (Ignoffo 1999) and public health settings (Cook et al. 2009) and has shown effectiveness in suppressing Australian mosquitoes, such as *Ae. vigilax* (Webb et al. 2012). An example of this approach is the In2Care® station, a mosquito contamination station designed to reduce local mosquito populations and potentially reduce the risk of disease transmission (https://www.in2care.org). The station lures in gravid container-breeding mosquitoes and contaminates them with the insect juvenile hormone analogue and insect growth regulator, pyriproxyfen, which affects larvae and pupae (Unlu et al. 2020). Contaminated mosquitoes spread the insecticide to other breeding sites so that mosquito larvae are not only controlled inside the station, but also in other (often cryptic) breeding sites in the vicinity.

Pyriproxyfen is combined with the entomopathogenic fungus *Beauveria bassiana* in In2Care® stations. Entomopathogenic fungi are necrotrophic parasites (i.e. kill their host in order to support their own growth and complete their life cycle), potentially useful for biocontrol (Lacey et al. 2015). *Beauveria bassiana* infects a broad range of hosts (Ferron 1978) such as ants (Broome et al. 1976), termites (Culliney and Grace 2000), agricultural pests such as grasshoppers (Bidochka and Khachatourians 1991) and arthropod vectors of human diseases, including mosquitoes (Howard et al. 2010, Bukhari et al. 2011). Infected mosquitoes usually die of the infection after a few days, reducing the length of time available to potentially transmit disease and removing mosquitoes from the breeding population. A reduction of mosquito density and effectiveness of the active ingredients of the In2Care® station have previously been validated by experiments on *Ae. aegypti* and *Ae. albopictus* (Buckner et al. 2017, 2021, Khater et al. 2022), but not for other *Aedes* species.

In this study, we aimed to evaluate the potential of the In2Care® station in reducing *Ae. notoscriptus* density in Victoria, Australia. We conducted a field trial using 110 In2Care® stations in a designated area and compared temporal changes in mosquito numbers to those in four control areas that did not receive the In2Care® station treatment. The efficacy of the In2Care® station was evaluated by comparing egg counts between all sites, as well as by measuring the larvicidal impact achieved in surrounding mosquito breeding sites due to pyriproxyfen autodissemination. We also conducted laboratory experiments to validate the effect of *B. bassiana* on adult mosquito survival. The results provide insights into the effectiveness of the In2Care® station in suppressing populations of *Ae. notoscriptus* and suggest a promising new approach for controlling this species in locations where it has become invasive.

## 2 MATERIAL AND METHODS

### 2.1 Field experiments

#### 2.1.1 Permits

This work was performed under APVMA Small-scale Trial Permit PER7250. Trials conducted to generate data relating to efficacy, residues, crop or animal safety or other scientific information outside the confines of a research facility where the size of the trial annually does not exceed the following: a total of 5 hectares nationally, with a maximum of 1 hectare in any one jurisdiction in the case of any food and/or fibre field crop (https://permits.apvma.gov.au/PER7250.PDF). The active ingredient In2Mix® sachets were imported under biosecurity permit 0006570560 from the Australian Government Department of Agriculture, Fisheries and Forestry. Permit conditions meant that the trial was limited to one treatment site, but we had access to several control sites.

#### 2.1.2 In2Care® stations

The In2Care® mosquito station is constructed of polyethylene and comprises several components, including a lid, central tube, detachable interface, and a reservoir filled with 3.5-4.5 litres of water infused with two yeast tablets to attract gravid mosquitoes. A floating platform moves up and down the central tube in response to the water level, providing a resting place for adult mosquitoes. This platform is equipped with a statically charged gauze strip coated with a powder containing pyriproxyfen and *B. bassiana*. When adult mosquitoes land on the gauze, the bioactives from the powder are transferred to them. The powder mixture, known as In2Care® Mix, is sealed in an aluminium refill sachet. Each sachet contains a 0.5 g powder formulation composed of approximately 74.03% pyriproxyfen and 10.00% *B. bassiana* strain GHA, with a minimum of 4.5 × 10^9 viable spores per gram. After depositing eggs, adult mosquitoes leave the station and disseminate pyriproxyfen to other breeding sources. Pyriproxyfen mainly inhibits the metamorphosis of mosquito pupae into adults and thereby prevents adult emergence. Exposure to the entomopathogenic fungus, *B. bassiana*, leads to mortality within 8-10 days. Station density of around one trap per 400 m^2^ is recommended, and traps require servicing every four to six weeks. During servicing, the powder-treated gauze strip is replaced, and the reservoir is refreshed with water, bioactive powder, and yeast tablets (http//www.in2care.org).

#### 2.1.3 Study sites

We assigned one treatment site and four control sites for this study: Eynesbury (In2Care® treatment) (Figure 1), Darley (C1), Aintree (C2), Diggers Rest (C3) and Wyndham Vale (C4). All sites were located north-west of Melbourne in Victoria, Australia and comprised ∼80 residential houses over a total ∼50,000 m^2^ area. Previous surveillance has shown that all sites have established populations of *Ae. notoscriptus* (Paris et al., unpublished). All sites represented residential areas with 1-2 storey houses on allotments of 572.42 m^2^ to 958.83 m^2^.

**Figure 1.**
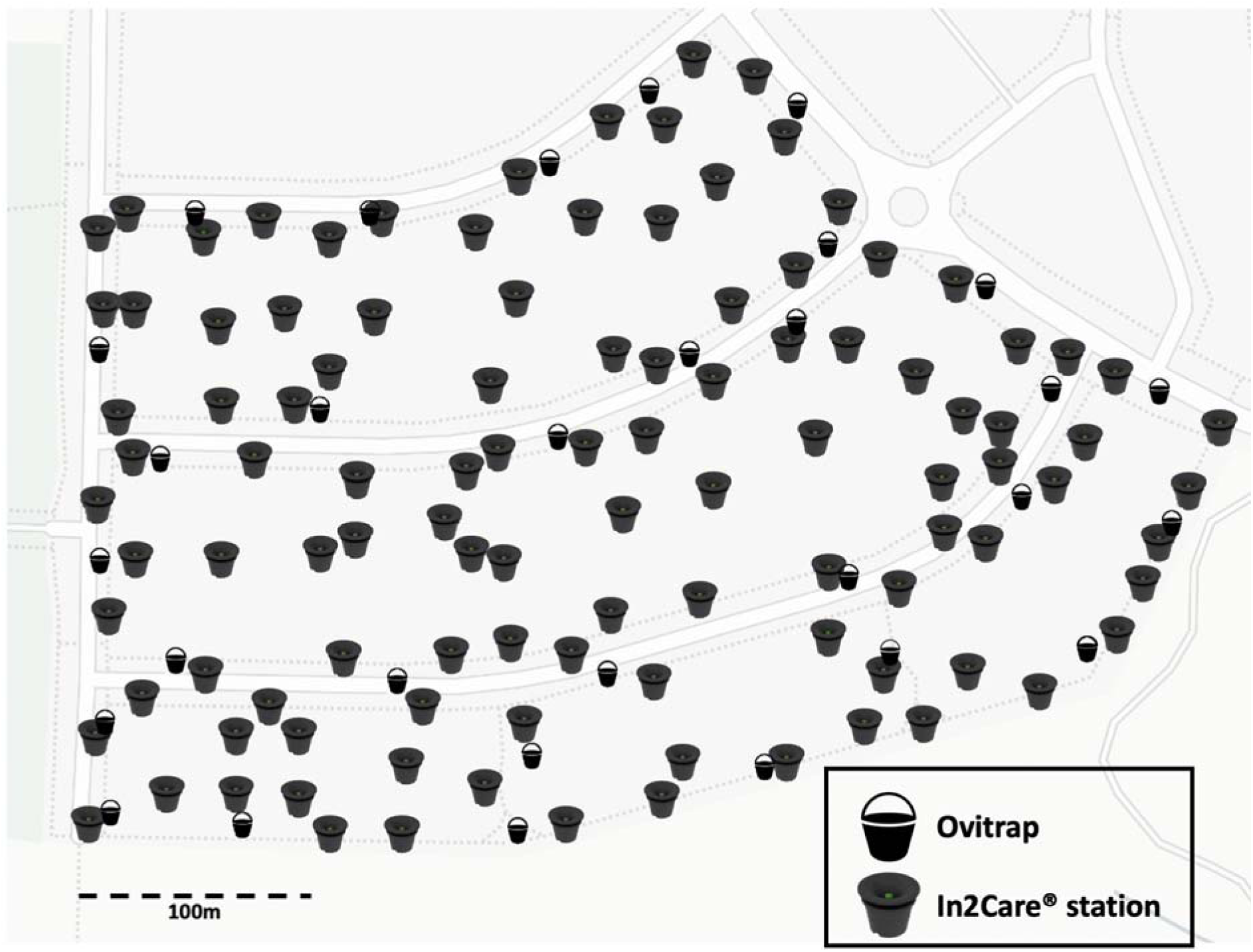
In2Care® station and ovitrap placement in the treatment site. Stations were placed at a density of one station per 450 m^2^, resulting in 110 stations deployed, represented by the In2Care® station symbol. Ovitraps (n=30) were placed on public land to monitor egg number before, during and after the intervention, shown by the bucket symbol (mean distance of ovitraps to stations = ∼158 m).

#### 2.1.4 Mosquito monitoring

Mosquito population density surveillance was conducted using egg counts at control and In2Care® treatment sites from the 20^th^ of January 2022 until the 21^st^ of April 2022. A total of 150 ovitraps (30 per site) were used, each consisting of a 500 mL black plastic bucket half-filled with water and containing alfalfa pellets to attract gravid *Ae. notoscriptus*. A 15 cm long strip of red felt extending into the water provided a consistent oviposition substrate. Egg counts were taken at weekly intervals for two weeks prior to the deployment of In2Care® stations. After the deployment of In2Care® stations, eggs were collected fortnightly, followed by weekly collections for two weeks after the intervention concluded. The same researcher performed all egg counts under a microscope to ensure consistency. To prevent the ovitraps from becoming functional breeding sites, all larvae and pupae present in the ovitraps were discarded at each collection.

#### 2.1.5 In2Care® station deployment, monitoring, and servicing

We placed In2Care® stations in volunteer properties, as well as on nature strips and other public land, with a density of approximately one station per 450 m^2^ close to the manufacturer’s recommendation (one station per 400 m^2^, https://www.In2Care.org). This resulted in a total deployment of 110 stations as shown in Figure 1. The stations were strategically located in shaded or semi-shaded areas that were expected to attract *Ae. notoscriptus*, such as areas with vegetation or proximity to surrounding breeding grounds. During the station deployment process, we filled the water reservoir with approximately 4.7 litres of tap water before attaching the netting containing the biocides to the floater and placing it onto the water surface. We aimed for a homogeneous distribution of the biocides by shaking each In2Mix® sachet containing the biocide powder, odour tablets, and netting well. The remaining biocide powder and odour tablets were placed into the water reservoir before the lid was locked into place.

Regular monitoring of the In2Care® stations was conducted on a fortnightly basis. The monitoring included checking for signs of disturbance, ensuring that the water level was sufficient, (i.e., filled more than halfway) and that the netting was still dry and intact. The presence of larvae and live and/or dead pupae in the stations was also recorded in order to validate the station’s attractiveness to gravid female mosquitoes and to ensure that no mosquitoes developed into adults inside the stations. When recording larval and pupal counts, we did not discriminate between *Ae. notoscriptus* and other species that might have been present in the stations. After four weeks, the biocides and odour tablets in the stations were refreshed with In2Mix® as previously described.

Weather data were obtained from the Bureau of Meteorology (bom.gov.au) Melbourne Airport Observation Station.

### Field study limitations

Our study had limitations that should be taken into consideration when interpreting the results.

Firstly, the study duration was relatively short, as the research permit was only approved after the start of the mosquito season and we were limited by the onset of autumn when mosquitoes become uncommon due to cold weather. Although this timeframe allowed for the intervention to be carried out for two mosquito generations, providing sufficient time to observe the effects of the stations, future interventions would benefit from an extended duration to capture a more comprehensive assessment of long-term effectiveness. Our research permit also limited us to a single treatment site. Future field studies will aim to incorporate multiple treatment sites to assess the scalability and general utuiliy of the approach.

In this study, population density was measured using egg counts. In *Ae. notoscriptus,* egg counts from ovitraps can correlate with genetic neighbourhood size, a parameter which varies proportionally with population density (Paris et al. 2023). In other *Aedes* species, ovitrap data can also be closely correlated with adult trap numbers across an environment (e.g. Tantowijoyo et al. 2016). Ovitraps have advantages over other types of traps, including their affordability, ready availability, and ease of servicing. These factors facilitate the deployment of a high number of traps to cover local heterogeneity in the environment.

### 2.2 Laboratory experiments

#### 2.2.1 Mosquito rearing

##### All laboratory experiments in this study used mosquitoes from laboratory colonies of

*Ae. notoscriptus* and *Ae. aegypti*. These colonies were established from field collections made in Cairns, Australia in 2019 for *Ae. aegypti*, and Brisbane, Australia in 2014 for *Ae. notoscriptus*. The colonies were maintained in a temperature-controlled insectary at 26°C ± 1°C, with a 12-hour photoperiod, as per the protocol outlined by Ross et al. (2017) for rearing of *Ae. aegypti*. The mosquitoes were provided with constant access to a 10% sucrose solution, and were housed in 27 cm^3^ BugDorm-1® cages (MegaView Science Co., Ltd., Taichung City, Taiwan). Adult females were blood-fed on a human volunteer’s arm, once per generation, to initiate egg-laying (with ethics approval from The University of Melbourne 0723847). The colonies were maintained in replicate cages, each containing approximately 500 mosquitoes.

#### 2.2.2 Laboratory validation of pyriproxyfen autodissemination

To determine whether pyriproxyfen was being disseminated from the In2Care® stations to other breeding sites, we collected fortnightly water samples from 15 of the ovitraps monitoring the treatment site. The traps were selected randomly after excluding ovitraps that did not contain any eggs or larvae which ensured selected ovitraps had been visited by mosquitoes. A total of 200 mL of water per trap was collected in a sealable plastic container which was then placed into an individual zip-lock bag to avoid cross-contamination between samples. We also collected 200 mL of water from five randomly selected In2Care® stations to serve as positive controls. Finally, five 200 mL water samples were taken from randomly chosen ovitraps from each control site as negative controls (20 samples total).

To measure residual intervention effects after the In2Care® stations were removed from the treatment site, we collected additional water samples from ovitraps in week 12 and 13 of the trial. We collected ten water samples from In2Care® stations in week 10 to act as positive controls for those additional samples.

All collected water samples were strained through fine mesh to remove any organisms or organic material that might have been collected with the water. We used individual pieces of mesh for each sample to avoid cross contamination between samples. We then added 25 L3 laboratory-reared *Ae. notoscriptus* larvae into each water sample and added two TetraMin® tropical fish food tables (Tetra, Melle, Germany). The containers were placed into a 26°C climate-controlled cabinet with a 12:12 light:dark cycle. All containers were checked daily to ensure larvae had sufficient food, and any mosquito eclosion was recorded. We used the number of free pupal exuviae as a proxy for successful adult eclosion.

#### 2.2.3 Laboratory validation of *B. bassiana* efficacy against *Ae. notoscriptus*

To validate the adulticidal effect of *B. bassiana* on adult *Ae. notoscriptus* and *Ae. aegypti,* we performed survival assays in the laboratory. All mosquitoes used in the experiment were adults seven days post-eclosion and were allowed to mate. We exposed 25 females to a fresh In2Mix® netting (containing approximately 0.2 g of In2Mix® powder with 74.03% pyriproxyfen and 10% *B. bassiana* spores) using the WHO standard forced exposure bioassay (The World Health Organisation 2006). Eight replicates of 25 females were prepared for the treatment and the control groups. We provided each replicate group with 10% sucrose solution. Individuals that died within one hour post experiment set up were excluded from all analyses. Sucrose solution was refreshed as needed. We scored survival daily until all individuals in each treatment cohort died. Individuals were scored as dead if no movement occurred after being manipulated with forceps. To test whether mosquitoes were still able to blood feed and produce eggs post infection, we blood fed groups on day two post exposure. We recorded the proportion of mosquitoes that successfully blood fed for each group and collected and counted eggs on day four and five of the experiment. We repeated the same setup with In2Mix® nets collected from In2Care® stations that were deployed in the field for four weeks.

We calculated the germination rate of spores used to infect the treatment groups to validate the spore quality and to ensure consistent spore qualities across replicates. Germination rate was also used to compare the quality of fresh In2Mix® with that of four-week-old nettings. To estimate germination rate, we inoculated three Sabouraud Dextrose Agar (SDA) plates for each used netting by rubbing a sterile cotton tip on the netting and transferring the spore sample onto the plates. The plates were then sealed with Parafilm® “M” Laboratory Film (Amcor Flexibles North America) and stored at 26°C for 24 hours. After incubation we identified germinated and not germinated spores (300 spores total) on three randomly chosen areas of each plate using a light microscope. The germination rate was calculated using the following formula: [number of germinated spores] ÷ [total number of spores counted (germinated + not-germinated)] x 100. We then averaged the calculated germination rate across the three replicate plates.

### 2.3 Statistical analyses

All statistical analyses were performed in R Studio v1.14 (R Core Team 2021). To investigate whether the egg counts differed by sample site and by sampling week, we used Generalized Linear Models (GLMER) with a Poisson distribution, using the package ‘*lme4’* (Bates et al. 2015). The sampling site and the sampling week were used as explanatory variables, along with an interaction term, and we added the specific trap as a random factor. We then selected the best fitting model by comparing the Akaike Information Criterion (AIC) of models including interactions to all nested models and the null model. The model returning the lowest AIC was used for analyses. We then compared variables using the ‘*posthoc’* package in R. We also followed a more conservative approach by undertaking one sample t-tests to determine whether the average numbers of eggs per trap in the treatment site differed from the control sites. These were run on the bi-weekly data as well as on combined data across the period when the treatment was expected to have a particularly large impact on mosquito numbers (weeks 6-10 after deployment).

To test whether sampling week significantly influenced the number of stations that were positive for larvae as well as having live or dead pupae, we fitted GLMERs for each variable, using sampling week as the explanatory variable and stations as a random factor. We tested whether the water samples collected from the treatment site ovitraps, the control sites ovitraps and the In2Care® stations differed in the percentage of mosquitoes that successfully eclosed by using Kruskal-Wallis tests (a Kolmogorov-Smirnov test revealed that the data were not normally distributed). We ran a Mantel test to determine whether the percentage of eclosed mosquitoes was correlated with the distance from the nearest In2Care® station.

To investigate whether mosquitoes infected with *B. bassiana* had a shorter lifespan than control groups, whether there were differences in survival between mosquitoes tested on fresh or older nettings, and whether there were differences in survival between *Ae. notoscriptus* and *Ae. aegypti*, we performed a Cox regression survival analysis using the ‘survival’ package in R (Therneau and T. Lumley 2015). The variables ‘species’, ‘treatment’ and ‘netting’ were used as explanatory variables, while the ‘replicate’ was added as a random factor. We ran independent sample t-tests to investigate differences in the proportion of mosquitoes blood feeding between control groups and groups infected with *B. bassiana*.

## 3. RESULTS

### 3.1 Mosquito surveillance

The results of our mosquito population density surveillance, which was performed by counting eggs in ovitraps, suggest that the number of eggs was not obviously linked to rainfall events, but did gradually decrease as conditions became colder (maximum weekly temperatures decreased from 32°C to 21°C and minimum weekly temperatures fell from 18°C to 12°C, Figure 2). The minimum temperature recorded during the trial was 11.9°C and the maximum temperature was 32.6°C. The GLMER analysis revealed that the number of eggs per trap was significantly influenced by the sampling site as well as the sampling week (z = 53.94, p < 0.001) . *Post-hoc* comparisons showed no significant difference in egg counts in control sites compared to the treatment site up to week 6 of the trial. At sampling week 8 (or 6 weeks after In2Care® stations were deployed) we found significantly lower numbers of eggs in the treatment site compared to C1 (p = 0.002) and C2 (p = 0.001). At sampling week 10 (8 weeks after station deployment), we found significantly fewer eggs collected in the treatment site compared to all control sites (C1: p < 0.001, C2: p < 0.001, C3: p = 0.003, C4: p = 0.003). In sampling week 12 (10 weeks after station deployment) there were significantly fewer eggs in the treatment site compared to C1 (p = 0.003), C2 (p = 0.001) and C3 (p < 0.001) and in sampling week 13 (11 weeks after station deployment) we found fewer eggs in the treatment site compared to C2 (p = 0.032) (Figure 3). We also confirmed, using t tests where data were pooled across traps within a site, that the average number of *Ae. notoscriptus* eggs was higher in the control sites than in the treatment site at sampling weeks 8, 10, 12 and 13 of the trial (p < 0.01 in each case) as well as in sampling week 13 (p = 0.019) as well as in the expected impact period in sampling weeks 8-12 combined (p < 0.001).

**Figure 2.**
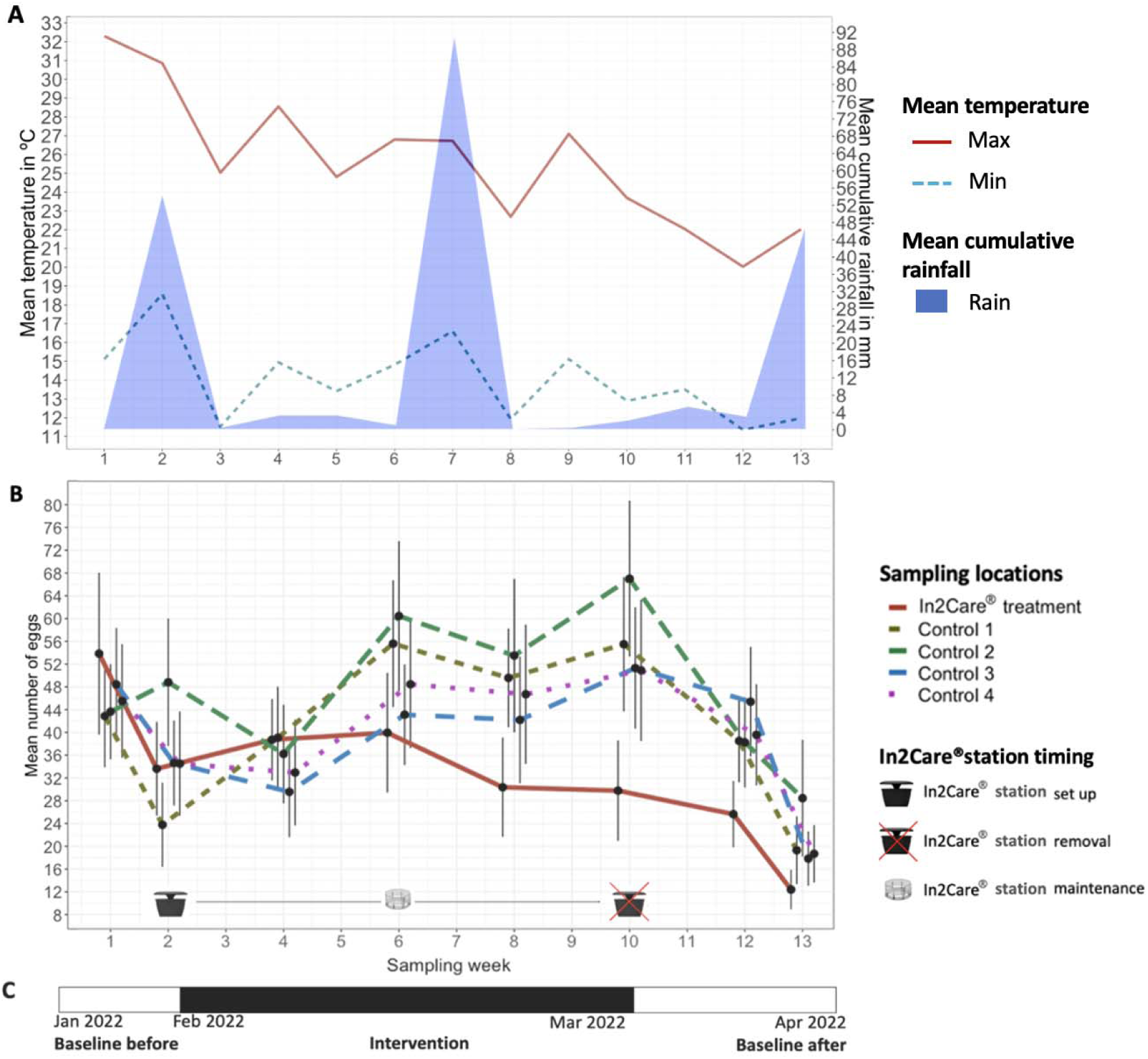
Egg counts and weather data during the In2Care® station trial. **(A)** Weekly mean minimum and maximum temperature are shown as red solid and blue double dashed lines respectively. Mean cumulative rainfall is represented by the light blue area plot. Weather data were obtained from the Bureau of Meteorology (bom.gov.au) Melbourne Airport Observation Station. **(B)** Weekly mean egg counts in the treatment site are shown by the solid red line. Dashed green, brown, purple, and blue lines represent egg counts in individual control sites. Egg counts are averaged from n=30 ovitraps per site. Black error bars show standard errors per collection and site. The In2Care® station symbols indicate the timing of station set and removal. The In2Care® floater symbol indicates maintenance of stations in week 6 of the trial (4 weeks post-deployment).

**Figure 3.**
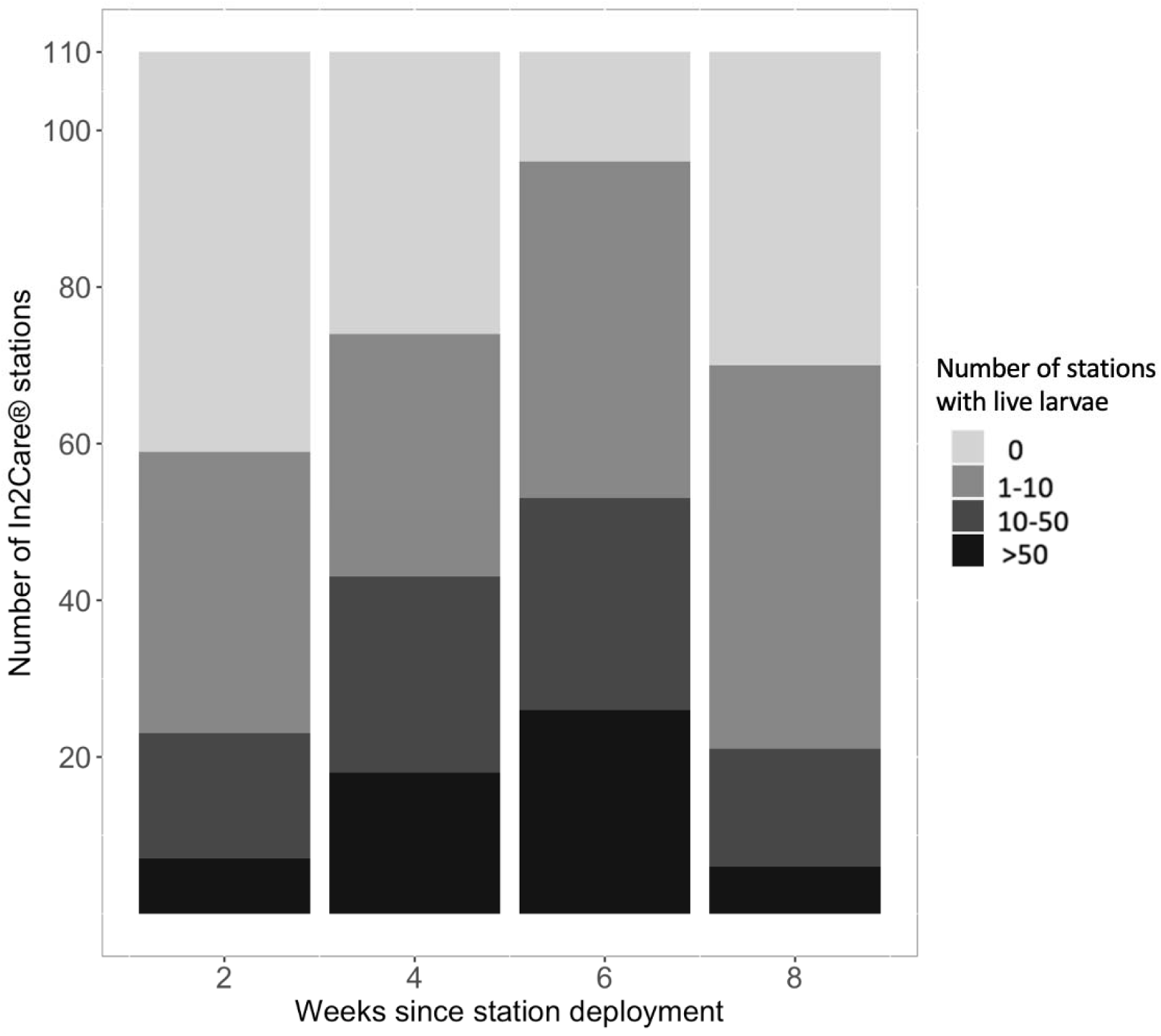
Number of In2Care® stations that presented live larvae since In2Care® stations were deployed. Number of larvae are represented by different shades of grey, with lighter shades indicating fewer larvae and darker shades more larvae n = 110 In2Care® stations per week. Fresh water and In2Mix® were added at 4 weeks.

Two weeks after the In2Care® stations were deployed in the treatment site, larvae were found in 53.64% (95% CI: 46.77% - 60.51%) of stations, indicating that at least half of stations had been visited by adult mosquitoes. After four weeks, 67.15% (95% CI: 58.22%, 76.32%) of stations contained immature mosquitoes. The number of stations containing immature mosquitoes increased to 87.27% (95% CI: 80.91% - 91.28%) after week 6 since deployment, and 63.63% (95% CI: 53.60% - 71.68%) after week 8 (Figure 3). Over the entire monitoring period, only two stations failed to show any signs of being visited; the reasons for this remain unknown. The GLMER showed that the weeks since initial deployment of the In2Care® stations significantly influenced the number of stations that were positive for larvae (GLMER: z = 23.87, p = 0.003). The numbers of live and dead pupae changed significantly across time (GLMER: alive: z = 12.68, p = 0.003; dead: z = 17.82, p < 0.002).

No In2Care® stations showed any pupae after being in the field for two weeks. After four weeks since In2Care® stations were deployed, 15.45% (95% CI: 14.96% - 15.95) of stations had live pupae in them. This increased to 81.82% (95% CI: 75.0% - 88.6%) after six weeks and 50% (95% CI: 40.43% - 59.57%) after eight weeks since deployment. Dead pupae were observed in In2Care® stations at six (48.18% (95% CI: 38.4% - 57.9%.) and eight weeks (89.09% (95% CI: 85.04% - 93.14%) since deployment (Figure 4).

**Figure 4.**
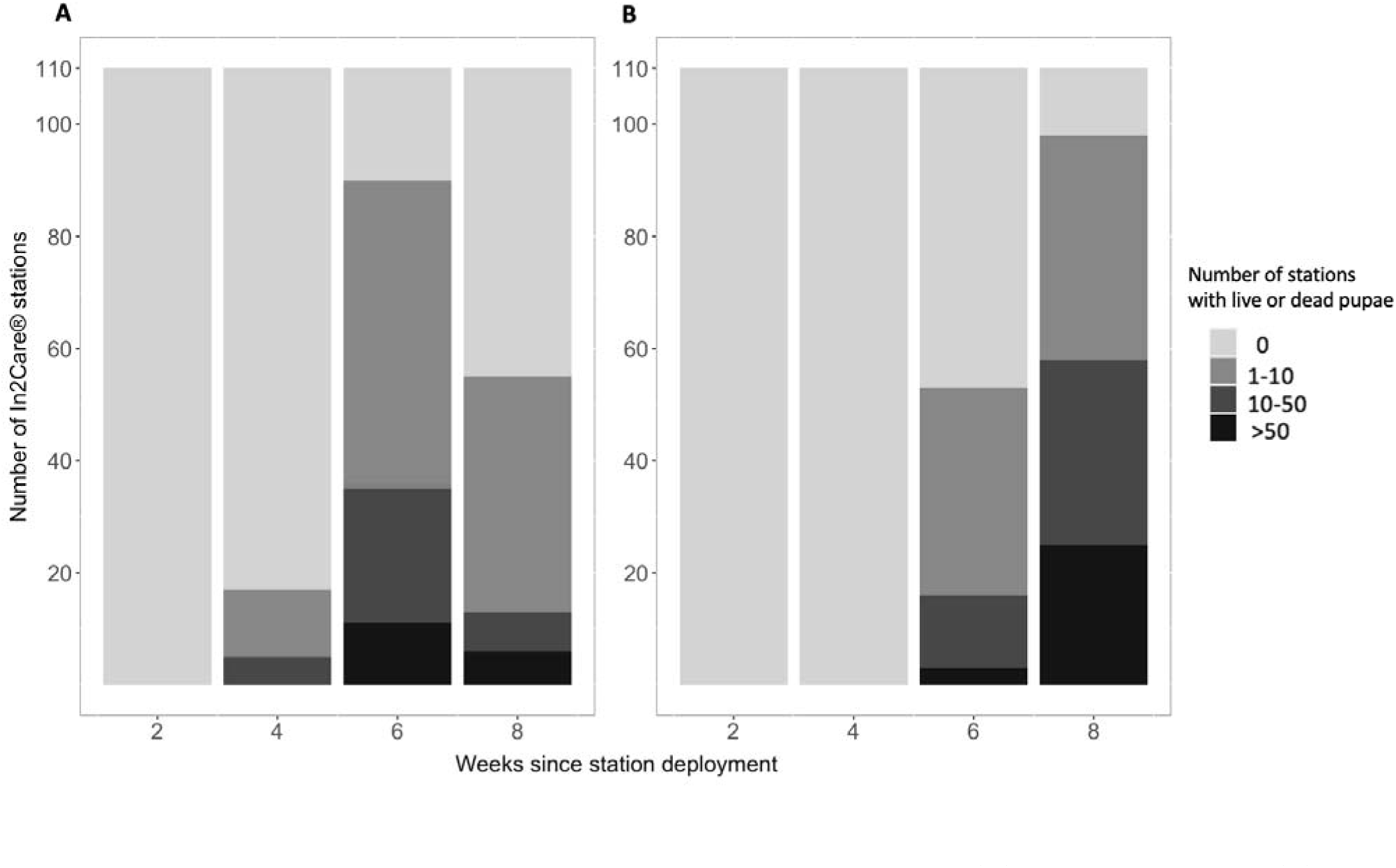
Number of In2Care® stations that presented live pupae (A) and dead pupae (B) since In2Care® stations were deployed. Number of pupae are represented by different shades of grey, with lighter shades indicating fewer pupae and darker shades more pupae n = 110 In2Care® stations per week.

### 3.2 Autodissemination

Significantly more pupae successfully eclosed when reared in water samples collected from control site ovitraps compared with water samples from In2Care® treatment site ovitraps (Kruskal-Wallis tests, X = -41.043, df = 2, p < 0.001) and In2Care® stations (X = 82.926, df = 2, p < 0.001). Eclosion of pupae in water samples collected from treatment site ovitraps were also significantly higher than samples collected from In2Care® stations (X = 41.925, df = 2, p < 0.001) (Figure 5). These results suggest that pyriproxyfen was autodisseminated from the In2Care® stations to the surrounding ovitraps. Baseline adult eclosion rates in the control site samples was on average 92%, compared to 42% in the intervention site.

**Figure 5.**
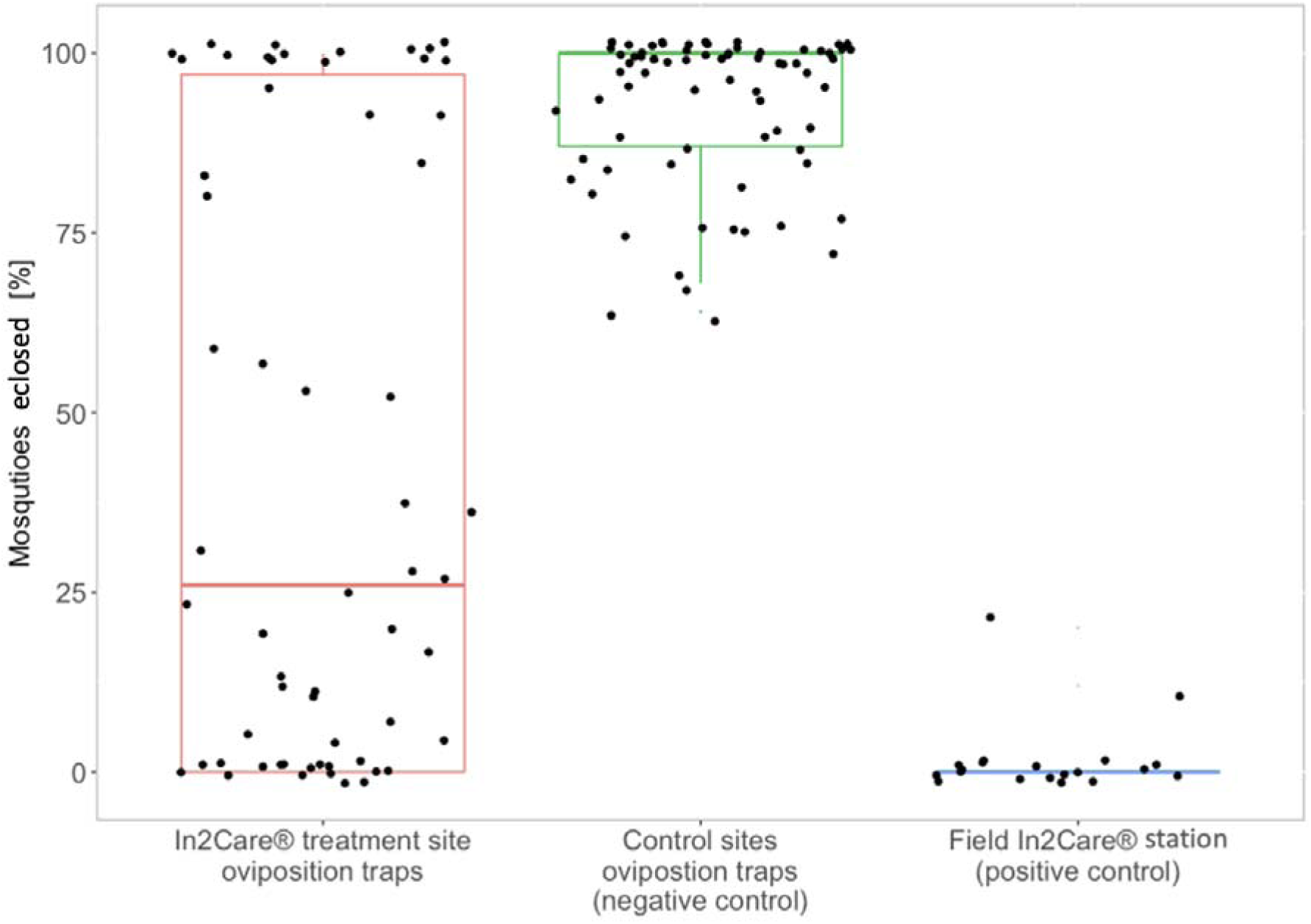
Percentage of eclosed mosquitoes from water samples collected during the In2Care® field trial. Eclosed mosquitoes from all collections combined. The red box represents water samples that were collected from ovitraps in the In2Care® treatment site (n=60). The green box represents water samples collected from control sites (n=64) and the blue box shows water samples collected from field In2Care® stations (n=20).

Autodissemination effects could also be observed from water samples collected from ovitraps two weeks after the In2Care® stations were removed. Aedes notoscriptus larvae had a significantly reduced percentage of eclosed adults compared to samples collected from control sites (Kruskal Wallis test: X = 15.83, df = 2, p < 0.001). Positive control samples (from In2Care® stations) showed 100% eclosion inhibition, which was significantly higher than the average 55% adult emergence inhibition in the treatment site ovitraps for week 12 and 13 combined (Kruskal Wallis test: X^2^= 26.380, df = 2, p < 0.001) (Figure 6), which also reflects the larvicidal effect of the In2Care® stations.

**Figure 6.**
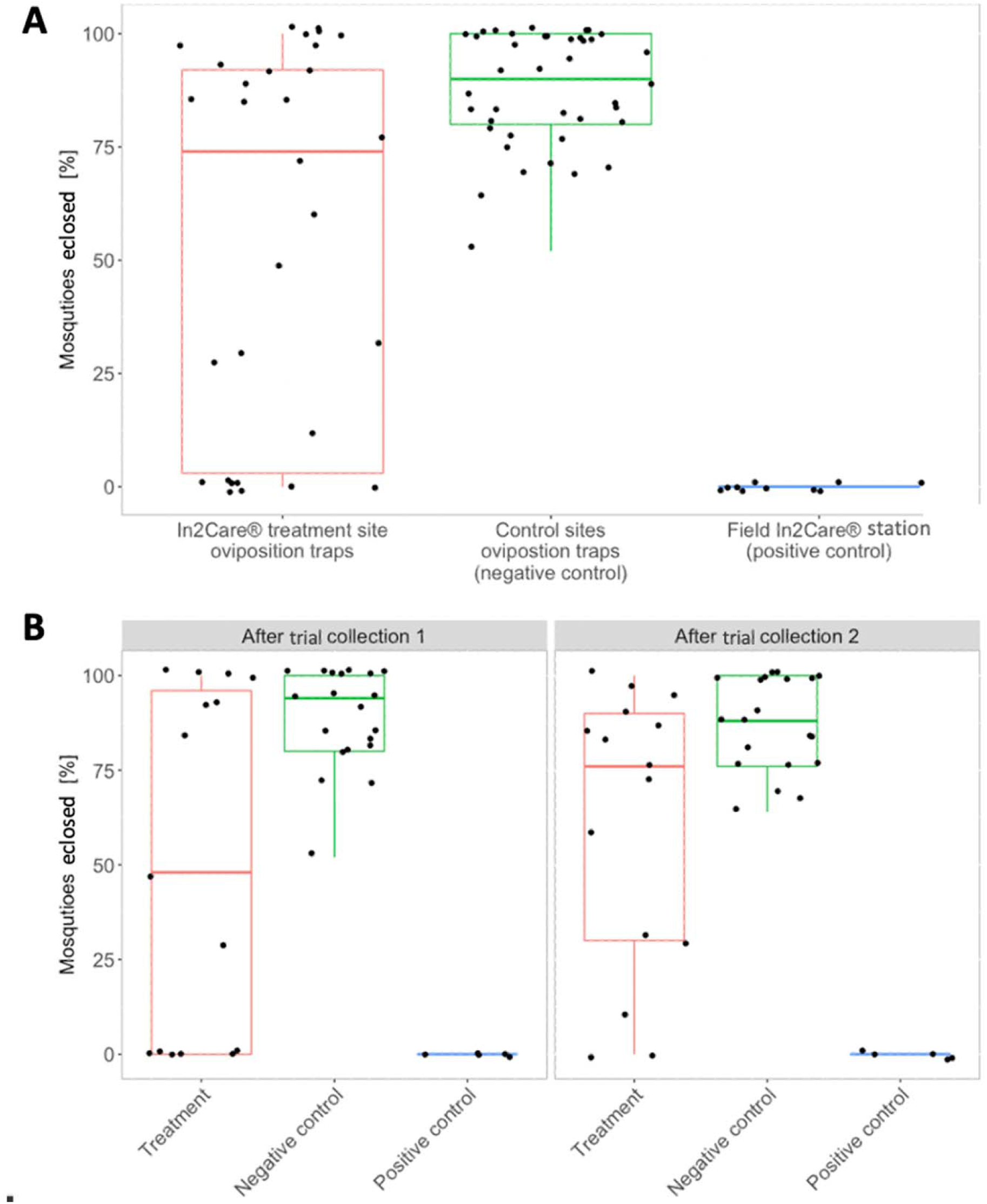
Percentage of eclosed mosquitoes from water samples collected after the In2Care® field trial. **(A)** Both collections combined for week 12 and 13 (2 and 3 weeks after removal of In2Care® Stations from the treatment site). Red boxes represent water samples that were collected from ovitraps in the In2Care® treatment site (n=30). Green boxes represent water samples collected from control sites (n=40) and blue boxes show water samples collected as positive control from field In2Care® stations in week 10 (n=10). Error bars indicate standard error. **(B)** Collections two and three weeks after removal of In2Care® Stations from the treatment site separately (In2Care® treatment site n=15; control sites n=20; In2care® stations n=5). Error bars indicate standard error.

We also tested whether the results differed between the week 12 and week 13 water collections to determine how long the residual effect of pyriproxyfen lasted in ovitraps. We found a significant difference in adult eclosion between the treatment site samples and samples from control sites (Kruskal Wallis test: X^2^ = 8.71, df = 2, p = 0.030) when collected two weeks after the removal of the In2Care® stations (week 12). The reductions in adult emergence inhibition were not significantly different in samples collected three weeks after station removal (Kruskal Wallis test: X^2^ = 7.92, df = 2, p = 0.260) although eclosion in controls was lower at this time (Figure 6). Baseline adult eclosing rates in the control site samples was on average 87%, compared to 55% in the intervention site.

There was no correlation between the proximity of the ovitraps to the closest In2Care® station and to all In2Care® stations (mean distance = ∼158 m; longest distance = ∼307 m; shortest distance = ∼4 m) and the percent of successful eclosion of mosquitoes from collected water samples (Mantel test: r = -0.05, p = 0.983; Figure S1).

### 3.3 Efficacy of B. bassiana on Ae. notoscriptus and Ae. aegypti

The results of the Cox-regression analysis indicated that there was a significant effect of the species tested (*Ae. notoscriptus* or *Ae. aegypti)* (z = -3.53, df = 3, p = 0.001), the age of the In2Care® netting (fresh or 4-weeks-old) (z = -59.49, df = 3, p < 0.001), and the treatment (infected or control) (z = 48.15, df = 3, p < 0.001) on survival. On average, survival of *Ae. aegypti* was 9% lower than *Ae. notoscriptus* (HR = 0.91 (0.87 – 0.96), p < 0.001). Both species had a four-times higher chance of dying if infected with In2Care® netting compared to uninfected control groups (HR = 4 (3.8 - 4.1), p < 0.001). If either species was infected using fresh In2Care® nettings, average survival was reduced by 88% compared to older nettings (HR = 0.12 (0.11 – 0.13), p < 0.001) (see Figure 7).

**Figure 7.**
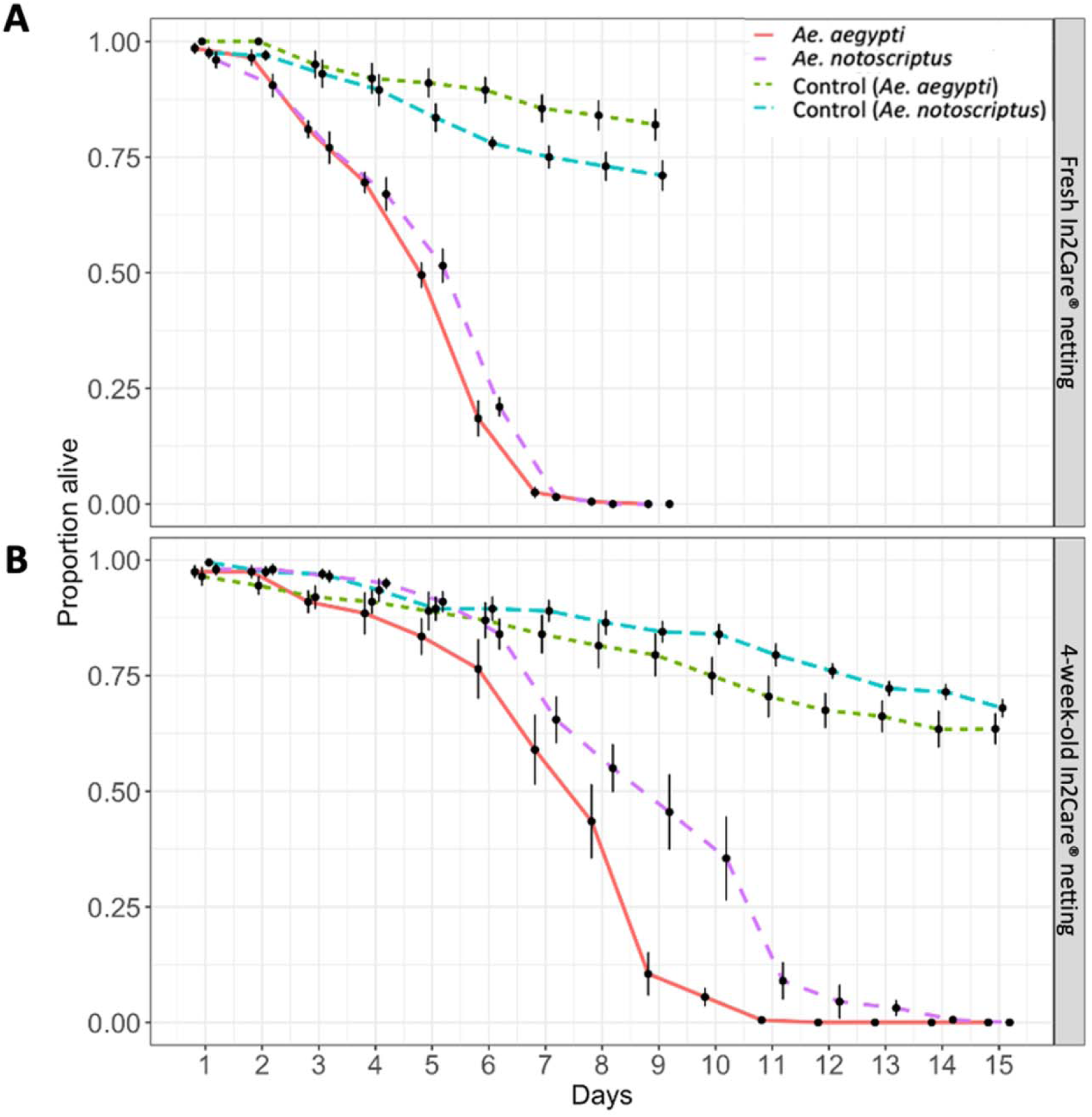
Survival of adult *Aedes notoscriptus* and *Aedes aegypti* after exposure to fresh In2Care® netting (A) and nettings that were in the field for four weeks (B). Solid lines represent the survival of the treatment groups with *Ae*. *notoscriptus* in magenta and *Ae*. *aegypti* in red. Dashed lines show the survival of control groups with *Ae*. *notoscriptus* in blue and *Ae*. *aegypti* in green. Data shows averaged proportion of survival from n = 8 replicates per group and species. Error bars indicate standard error.

We calculated an average Beauveria spore germination rate of 93.41% (95% CI: 91.18% - 97.79) for the fresh In2Mix® samples with no replicate being lower than 90%. The four-week-old nettings presented an average germination rate of 53.74% (95% CI: 49.92% - 57.71%).

We did not find a significant difference in the proportion of females that blood fed, between the infected and control groups (t-test: fresh In2Care® mix: Ae. notoscriptus: t = -0.03, df = 7, p = 0.98; Ae. aegypti: t = -0.01, df = 7, p = 0.41; 4-weeks old In2Care® mix: Ae. notoscriptus: t = 0.88, df = 7, p = 0.79; Ae. aegypti: t = 0-.34, df = 7, p = 0.74) (Figure 8). We did not observe oviposition in any of the infected groups, while control groups laid eggs.

**Figure 8.**
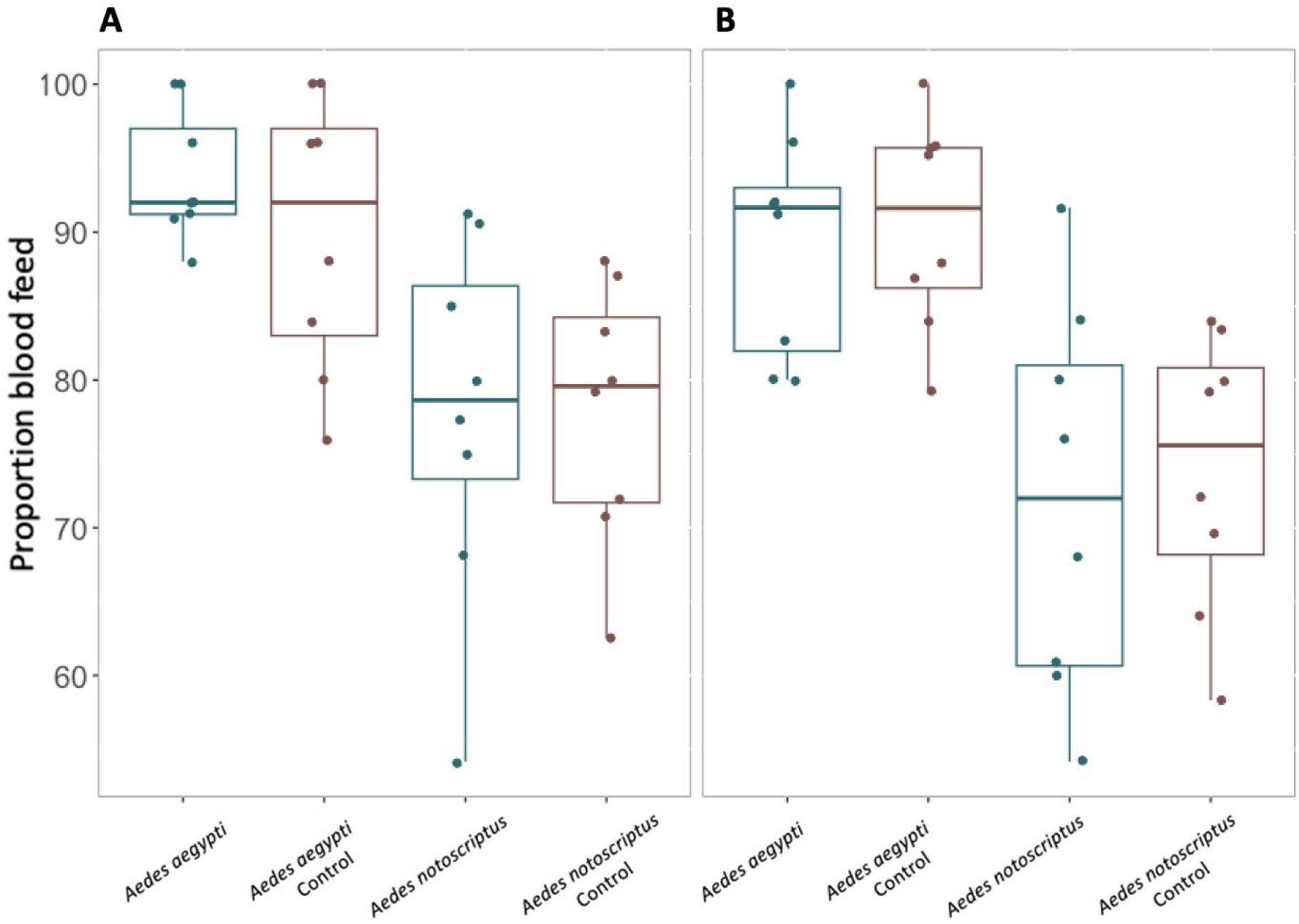
Proportion of mosquitoes that blood feed after B. bassiana infection. **(A)** Fresh In2Care® netting **(B)** 4-weeks-old In2Care® netting. Blue boxes indicate the proportion that blood feed of the treatment groups, red boxes indicate the proportion that blood feed in the control groups. Data is averaged from n = 8 replicates per group and species. Error bars show standard error.

## DISCUSSION

The results from our study suggest that the In2Care® station can effectively control the container breeding mosquito *Ae. notoscriptus*. We found a reduction in egg numbers in the treatment site after In2Care® stations had been deployed for six weeks with the effect persisting until three weeks after station removal. The pyriproxyfen autodissemination results indicate that the In2Care® stations were visited by mosquitoes which then successfully contaminated surrounding ovitraps with pyriproxyfen and that the disseminated larvicide persisted at lethal levels for at least two weeks after the In2Care® stations were removed. Additionally, we found that the fungal pathogen *B. bassiana*, which is incorporated in the In2Care® station, significantly decreased the lifespan of mosquitoes in a laboratory setting.

The In2Care® stations were successful in attracting mosquitoes, as shown by the high proportion of stations that housed larvae (Figure 3). After two weeks of deployment, slightly more than half of the stations contained larvae. This trend continued to increase over following weeks, before dropping somewhat to 63% after eight weeks. The decline in positive stations could indicate a decrease in the local *Ae. notoscriptus* population, especially considering that the traps were serviced at the 4-week mark, ensuring that the attraction levels of the stations for mosquitoes remained similar to those observed at the beginning of the intervention. A lower number of new eggs laid in the stations, as well as continued development of existing larvae to the pupal stage, would also have contributed to a decrease in larval numbers. The reduction of larvae in stations is further supported by an increase in the number of pupae after four weeks (Figure 4). The reduction in live pupae is likely caused by pyriproxyfen-induced mortality, which was observed in approximately half of the stations at six weeks post-deployment and in 87% of stations after eight weeks of deployment. Pupal mortality may have begun earlier than detected, as larvae in the stations may have fed on dead pupae. These findings also suggest that the In2Care® stations function as an ’egg dump’ that contributes to the overall reduction of the mosquito population, as eggs laid in the stations do not mature into adult mosquitoes.

Mosquito population sizes naturally fluctuate with rainfall and temperature (Bomblies 2012), with rainfall triggering egg hatching and higher temperatures accelerating development. We found that the general trend of egg counts in all sites followed these weather parameters (Figure 2). However, after six weeks of the In2Care® stations being in place, we observed a significant reduction in egg counts in the treatment site (Figure 2). The decrease in egg numbers at the treatment site cannot be attributed to weather factors alone, as all control sites displayed increasing egg numbers, while the treatment site showed a decline (Figure 2).

Our experiments on water collected from ovitraps at the treatment site confirm the effective autodissemination of pyriproxyfen. The results reveal a significant average decrease of about 50% in eclosing of adults in water samples from the treatment site compared to control sites (Figure 5). This finding suggests that mosquitoes carried pyriproxyfen from the In2Care® stations to ovitraps and likely to other surrounding breeding sites. Female *Aedes* mosquitoes tend to distribute their eggs across multiple breeding sites instead of laying them in one place, which helps reduce the risk of losing all offspring if the chosen breeding site proves to be unfavourable (Colton et al. 2003). This strategy of ‘skip oviposition’ favours the autodissemination of pyriproxyfen, as it increases the number of breeding sites that mosquitoes use. The autodisseminated pyriproxyfen likely contributed to the decrease in mosquitoes in the treatment site, with a cumulative effect over multiple generations. Importantly, the positive impact of pyriproxyfen persisted for two weeks even after the removal of the In2Care® stations from the treatment site (Figure 6). Note that we did not quantify the presence of any other mosquito species using the stations or ovitraps in our trial. Therefore, we cannot rule out the possibility that other mosquito species may have also contributed to the autodissemination of pyriproxyfen.

The potential for mosquitoes to distribute pyriproxyfen to other parts of the ecosystem where it would influence other insect systems should be considered before widespread application of the technology is implemented, as some laboratory studies have shown pyriproxyfen transfer by male mosquitoes (artificially dosed with high quantities of pyriproxyfen) to bees during nectar feeding (Kancharlapalli et al. 2021). However, pyriproxyfen has already been registered and approved globally (including in Australia) for widescale agricultural applications given its acceptable risk for non-target organisms and pollinators. If studies confirm these effects are found to be minimal or less damaging than the effects of broadscale mosquito control methods, then implementation of the In2Care® station will be of benefit in reducing the numbers of *Ae*. *notoscriptus* and other container-breeding mosquitoes.

We tested the adulticidal effect of *B. bassiana* on *Ae. notoscriptus* and compared the adulticidal effect of *Ae. notoscriptus* to *Ae. aegypti*, using both fresh In2Mix® and nettings that had been in the field for four weeks. Both species had a 75% higher chance of reduced survival if infected with In2Mix®-treated netting compared to control groups, and using fresh In2Care® nettings significantly reduced the average survival rate of both species by 88% compared to older nettings (Figure 7). These results show that older nettings still infect and kill adult *Aedes* mosquitoes, though their lifespan increases by approximately three to six days. This indicated that the nettings should be refreshed after four to six weeks, as recommended by the manufacturer. Our laboratory investigations suggest that *B. bassiana* used in the In2Care® technology has the potential to be an effective tool in controlling mosquito populations and has most likely contributed to the reduced egg densities found in this field intervention.

Our results showing a reduction in mosquito density as well as the effect of pyriproxyfen and *B. bassiana* are consistent with studies investigating the effect of the In2Care® station on *Ae. aegypti, Ae. albopictus* and *Culex quinquefasciatus* (Buckner *et al*. 2017; Buckner *et al*. 2021; Khater *et al*. 2022; Su et al. 2021) and suggest that the In2Care® stations successfully kill larvae of container breeding mosquitoes by spreading pyriproxyfen to surrounding breeding sites. The In2Care® station, in conjunction with source reduction and other integrated vector management tools, has potential to provide vector control and reduce the density of mosquito populations which would be expected to reduce the transmission of *Aedes*-borne arboviruses, parasites, and mechanically transmitted bacteria. Benefits of the In2Care® station to public health may extend beyond the reduction of mosquito vectors. Broadscale insecticide application may be obviated in an In2Care®-treated area, thereby reducing public exposure as well as off-target environmental impacts.

Though the results present a promising demonstration of the potential for the In2Care® station for *Ae. notoscriptus* control, there may be challenges in implementing the approach in some areas. Firstly, the cost of the stations is higher compared to traps used for similar purposes such as gravid traps, which may limit the cost-effectiveness of a large-scale rollout. Prices can vary between countries, and costs may decrease as stations become more widely used. However, the stations can be maintained over a long period of time, with only the active ingredients needing replacement, resulting in relatively low maintenance costs. Furthermore, the design of the stations is user-friendly, allowing households to maintain the traps. This feature reduces reliance on health authorities for trap deployment and maintenance, making interventions through community-based efforts more feasible.

In addition, in our trial, the treatment and control sites were in an isolated location, reducing the potential for migration of mosquitoes back into the treated area. Implementation of the strategy in less isolated areas may require the treatment of larger areas. Our study took place in a new housing development, where most houses are occupied and there are few empty blocks or construction sites, making it easier to place traps. We did identify a few mosquito breeding containers (mostly bird baths or drip trays of pot plants), but the area is generally well maintained, reducing the incidence of cryptic breeding sites. In areas with a low occupation rate of houses, there may be an increased risk of overflowing gutters and containers in backyards that can act as breeding sites. It is worth noting that in a study utilising the In2Care® station conducted in Hawaii (Brisco et al. 2023), a failure to find a reduction in mosquito numbers was partly attributed to numerous cryptic breeding sites in properties that did not participate in the intervention (in addition to an inadequate number of In2Care® stations being deployed), and source reduction measures may be required in some contexts.

## CONCLUSION

The In2Care® station was found to be effective in controlling a population of the container breeding mosquito *Ae. notoscriptus* in Victoria, Australia. The station demonstrated a reduction in egg numbers in the treatment site after six weeks of deployment. The autodissemination results indicate that the In2Care® stations were visited by mosquitoes which then successfully contaminated ovitraps with pyriproxyfen, and the larvicidal effect persisted for at least three weeks after the In2Care® stations were removed. The experiments validated the adulticide effect of the fungal pathogen *B. bassiana* on *Ae. notoscriptus* and *Ae. aegypti* and suggest that it has the potential to be an effective tool in controlling mosquito populations.

## ACKNOWLEDGEMENTS

Authors thank the Melton Council, especially Lawrie Conole and residents for permission and assistance. Jessica Home, Sophie Collier, Matthew Whitney, and Sarah Jaboor are thanked for assistance with fieldwork and lab assays. We thank Tim Möhlmann and Marit Farenhorst of In2Care Trading B.V., and Steve Broadbent of Ensystex for assistance in planning the intervention, lab experiments, provision of In2Care® stations and In2Mix® refills used in this study and for feedback on the manuscript.

## FUNDING

The study was funded by the National Health and Medical Research Council Partnership Project grant 1196396 ‘Stopping Buruli ulcer in Victoria’. VP was financially supported by the Australian Government Research Training Program Scholarship.

## Notes

### Competing Interest Statement

The authors have declared no competing interest.

### Summary of Updates

- New section explaining the In2Care stations - New section discussing limitations of field intervention - Updated statistics - Minor changes throughout

