## Supplementary material for "Evaluation of In2Care® mosquito stations for suppression of the Australian backyard mosquito, *Aedes notoscriptus*": Figure S1

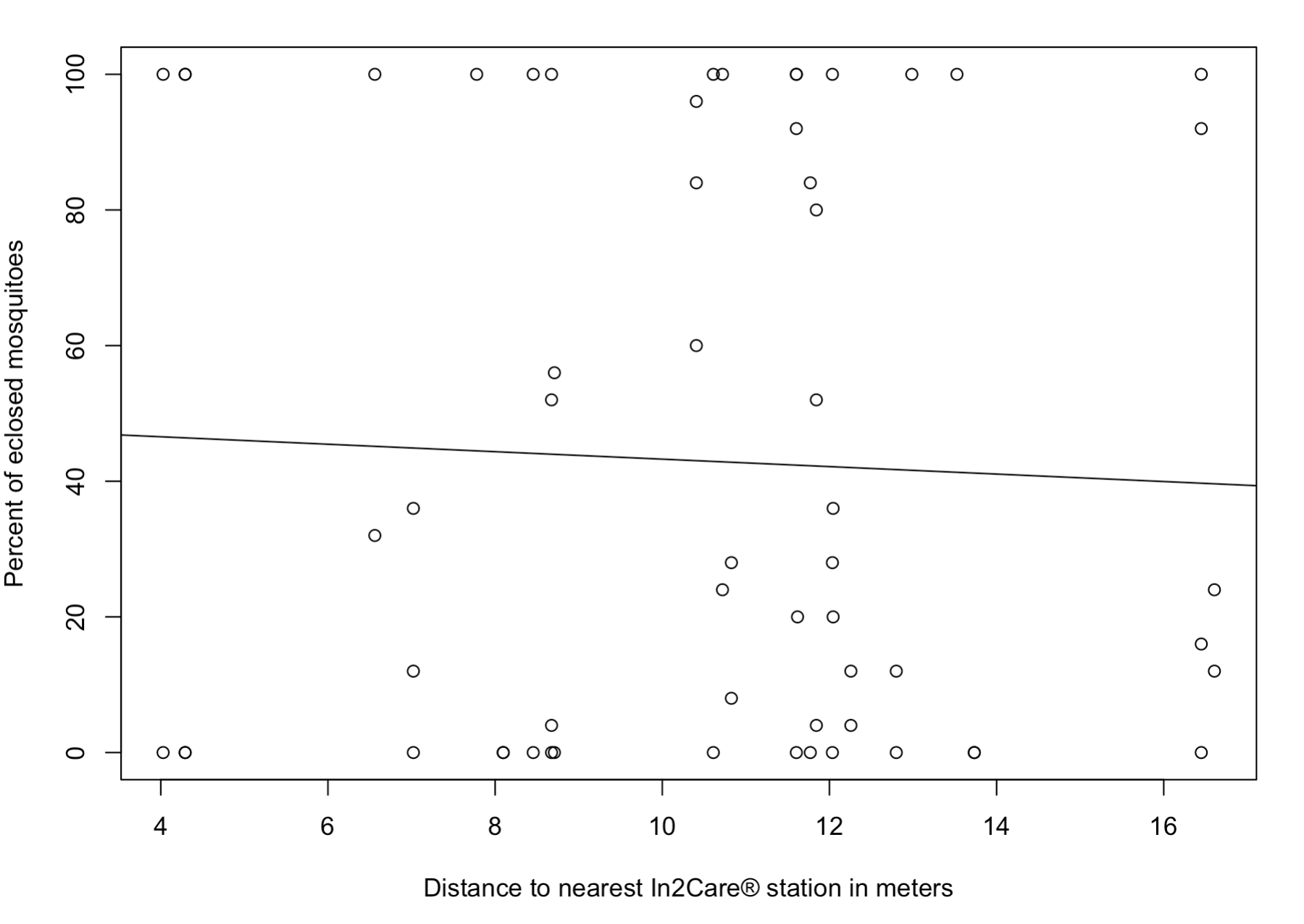

**Figure S1 Scatter plot of a Mantel test between the distance of water samples collected from ovitraps in the treatment site to the nearest In2Care® station (x-axis) and the percent of eclosed mosquitoes from those water samples (y-axis).** Line describes linear regression fit.
